# Similarities and differences between brain and skin *GNAQ* p.R183Q driven capillary malformations

**DOI:** 10.1101/2024.06.19.599711

**Authors:** Sana Nasim, Colette Bichsel, Anna Pinto, Sanda Alexandrescu, Harry Kozakewich, Joyce Bischoff

**Author notes:** Corresponding authors:* Joyce Bischoff, PhD, Karp Family Research Laboratories 12.212, Boston Children’s Hospital, 300 Longwood Ave., Boston, MA 02115, Sana Nasim, PhD, Karp Family Research Laboratories RB12004A, Boston Children’s Hospital, 300 Longwood Ave., Boston, MA 02115.

## Abstract

Capillary malformations (CM) are congenital vascular irregularities of capillary and venous blood vessels that appear in the skin, leptomeninges of the brain, and the choroid of the eye in the disorder known as Sturge Weber Syndrome (SWS). More common are non-syndromic CM found only in the skin, without brain or ocular involvement. A somatic activating mutation in *GNAQ* (p.R183Q) is found in ∼90% of syndromic and non-syndromic CM specimens and is present in CD31^pos^ endothelial cells isolated from brain and skin CM specimens. Endothelial expression of the *GNAQ* p.R183Q variant is sufficient to form CM-like vessels in mice. Given the distinct features and functions of blood vessels in the brain versus the skin, we examined the features of CM vessels in both tissues to gain insights into the pathogenesis of CM. Herein, we present morphologic characteristics of CM observed in specimen from brain and skin. The *GNAQ* p.R183Q variant allelic frequency in each specimen was determined by droplet digital PCR. Sections were stained for endothelial cells, tight junctions, mural cells, and macrophages to assess the endothelium as well as perivascular constituents. CM blood vessels in brain and skin were enlarged, exhibited fibrin leakage and reduced zona occludin-1, and were surrounded by MRC1^pos^/LYVE1^pos^ macrophages. In contrast, the CMs from brain and skin differ in endothelial sprouting activity and localization of mural cells. These characteristics might be helpful in the development of targeted and/or tissue specific therapies to prevent or reverse non-syndromic and syndromic CM.

**Statements and Declarations:** None

## Introduction

Sturge Weber Syndrome (SWS) is a non-inherited, rare neurocutaneous disorder with atypical morphology and function of capillary and venous blood vessels. SWS is strongly associated with a cutaneous capillary malformation (CM), also known as port-wine birthmark, leptomeningeal vascular malformation in the brain, visible with contrast enhanced magnetic resonance imaging (MRI), and increased intraocular pressure and choroidal ‘hemangioma’[1, 2]. Some of the neurological manifestations include seizures, stroke-like episodes, hemiparesis, cerebral atrophy, calcification, developmental delays, and cognitive limitations. Patients with both upper and lower eyelid port wine birthmark are at high risk of developing glaucoma, which can lead to optical nerve damage and loss of vision[2]. Currently, there are no molecularly targeted drug therapies to treat SWS. Some patients, considered non-syndromic, have cutaneous CMs but no brain or ocular involvement.

As seen by histopathology, CM is composed of dilated, enlarged vessels with irregular shapes and thickened basement membranes[3]. Furthermore, CM vessels can be surrounded by multiple layers of disorganized mural cells or lack mural cell coverage altogether[3, 4]. CM endothelial cells (ECs) in non-syndromic cutaneous lesions appear to be immature as they express stem cell markers (CD133, CD166) as well as venous (EphrinB1) and arterial (EphrinB2) markers[3]. CM ECs in the leptomeninges show strong expression of HIF1α and HIF2α, which suggests sustained angiogenic signaling[5]. We recently showed strong expression of intracellular adhesion molecule-1 (ICAM-1) in ECs in leptomeningeal CM in SWS brain specimens and increased MRC1^pos^/LYVE1^pos^/CD68^pos^ macrophages; such cells were not found in non-SWS brain[4].

Syndromic and non-syndromic CM are caused by a somatic activating mutation in *GNAQ*[6], the gene that encodes G protein subunit alpha-q (Gα_q_). The *GNAQ* p.R183Q mutation, located in the switch 1 domain of Gα_q_, causes constitutive activation of Gα_q_. The mutation is found in ∼90% of port wine birthmark and ∼88% of brain lesions from SWS patients[6]. Recent testing of an additional patient cohort identified mutations in the *GNAQ* homolog *GNA11* and in *GNB2* which encodes the G protein subunit beta 2, a protein that forms a heterotrimeric complex with Gα_q_[7, 8]. The *GNAQ* mutation is highly enriched in EC isolated from brain and skin CM specimens, yet there are indications that other unknown cell types may carry the p.R183Q mutation[9-11].

Our motivation here was to assess differences between the CM in brain and skin by comparing histopathological characteristics. Our ultimate goal was to understand how the single amino acid change in Gα_q_ disrupts capillary-venous morphology in these two distinct tissues. Such insights could help in the development of targeted molecular and tissue-directed therapies to alleviate the morbidity and social impacts of CM and SWS.

## Materials and Methods

### Reagents and antibodies

#### Primary antibodies

Alpha smooth muscle actin Cy3 conjugated (mouse, 1:500, Sigma Aldrich, C6198); Calponin (rabbit, 1:100, abcam, ab46794); Desmin (rabbit, 1:100, abcam, ab15200); LYVE1 (goat, 1:150, R&D systems, AF2089SP); MRC1 (mouse, 1:100, Millipore Sigma, AMAB90746); Zona occludin-1 (mouse, Thermo Fisher, 33-9100).

#### Secondary antibodies

Goat anti-Rabbit IgG (H+L), Alexa Fluor™ 488 (1:200, Invitrogen, A11008); Donkey anti-Mouse IgG (H+L), Alexa Fluor™ 488 (1:200, Invitrogen, A11008) Donkey anti-Goat IgG (H+L), Alexa Fluor™ 594 (1:200, Invitrogen, A32758).

#### Other reagents

4’,6-diamidino-2-phenylindole - DAPI (Invitrogen, R37606); Ulex europaeus agglutinin-I, UEAI (1:100, Vector labs, RL-1062)

### Droplet digital polymerase chain reaction

Human brain and skin specimens (**Table 1**) were obtained under a human subject protocol (IRB-P00003505) approved by the Committee on Clinical Investigation at the Boston Children’s Hospital. Droplet digital PCR (ddPCR) was used to measure the mutant allelic frequency in tissue sections as previously described [12]. Briefly, DNA was isolated from formalin-fixed paraffin embedded (FFPE) tissue sections using a FormaPure kit (Beckman Coulter, C16675). ddPCR with probes to detect the *GNAQ* R183Q mutation (**Table S1**) was run with 15ng DNA per sample along with ddPCR SuperMix (BioRad, 1863010). The droplet fluorescence was read in a QX200 droplet reader and analyzed in QuantaSoft software (BioRad).

**Table 1.**
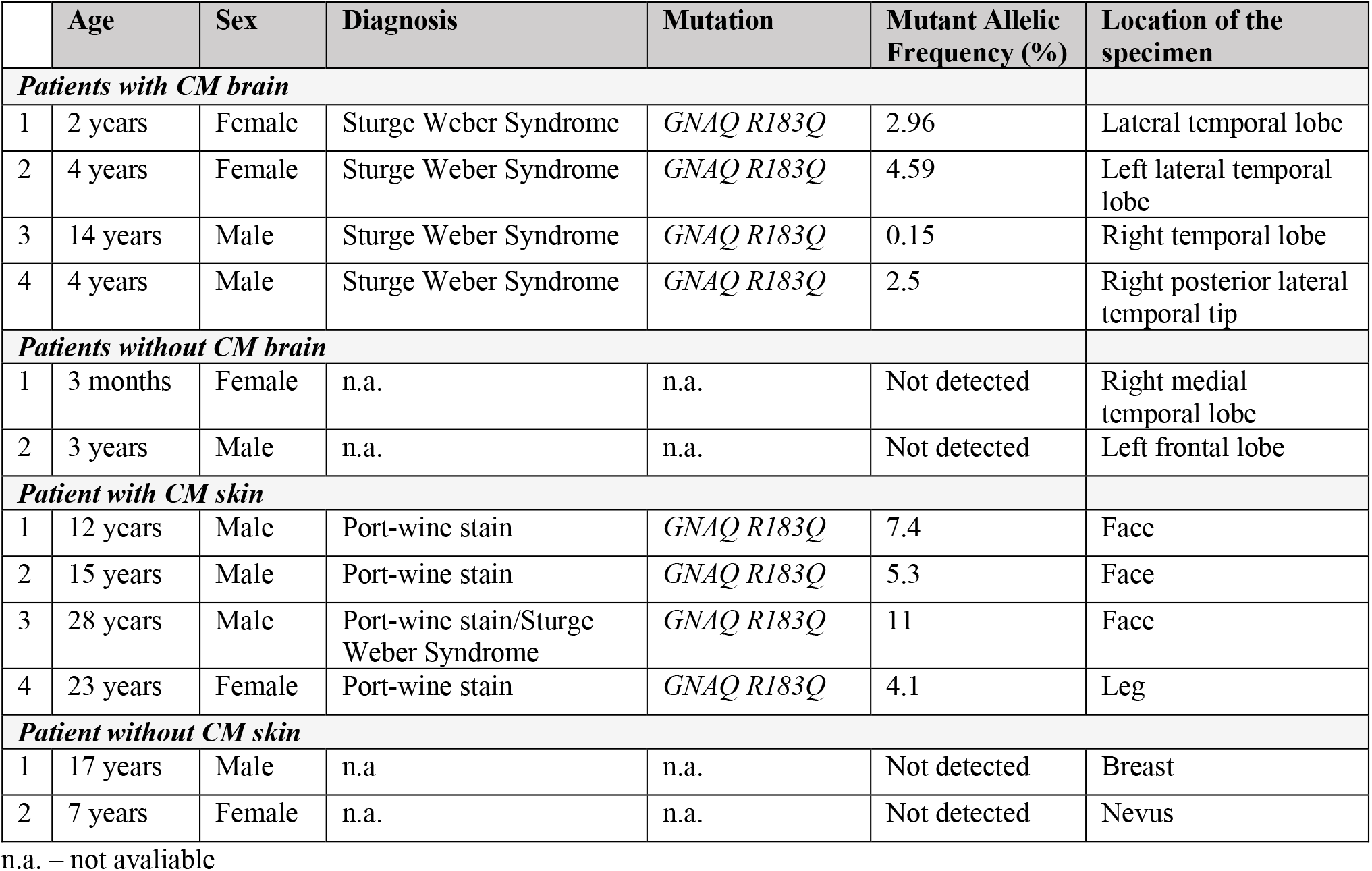
Patient demographic information for brain specimens.

### Immunofluorescence Staining

5*μ*m sectioning of human brain and skin specimens (**Table 1**) was performed by the Boston Children’s Hospital Department of Pathology. Sections were deparaffinized in xylene and series of ethanol incubation followed by 1X universal HIER antigen retrieval (abcam, ab208572) for 20 mins at 95°C. Sections were incubated in blocking buffer (10% normal goat serum (abcam, ab7481), 0.3% Triton X-100, 1% bovine serum albumin (Millipore Sigma, A8806) in phosphatase buffer saline (PBS)) at room temperature for 1 hour followed by primary antibody incubation at room temperature for 1 hour or at 4°C overnight. The primary antibody was diluted in a buffer containing 1% normal goat serum, 0.3% TritonX-100, and 1% bovine serum albumin. Sections were thoroughly washed (1X PBS) before and after incubating in secondary antibody for 1 hour at room temperature. Lastly, slides were counterstained for 4′,6-diamidino-2-phenylindole (DAPI) (Invitrogen, R37606) for 5-10 mins before mounting. Images were taken on Zeiss Laser Scanning 880 confocal microscope and analyzed in Fiji software.

### Image Quantification

Vessels with sprouting UEAI^pos^ cells were quantified manually. Blood vessels with one or more ECs that appeared to be sprouting away from the vessel lumen were counted as a sprouting vessel. For quantification, the data was normalized to total number of vessels/mm^2^. For mural cell quantification, all the vessels were manually categorized for three unique features: mural cell layer, no mural cell layer and distant mural cell layer. The quantification of distance between the calponin-positive or desmin positive vessels from the UEAI^pos^ endothelium was also measured. MRC1 and MRC1/LYVE1 colocalization were quantified as previously described[4]. In brief, for automatic colocalization, the Pearson Correlation Coefficient (PCC) and the Manders Overlap Coefficients served as the primary indicators for fluorescent colocalization. The PCC serves to represent how closely two markers follow a simple linear relationship. The Manders Overlap Coefficients--denoted by tM1 and tM2--reveal the proportion of cell area that shows colocalization of markers in comparison to the total measured fluorescence of one marker[13]. Next, the Costes Method of colocalization analysis was applied using the Coloc2 plugin in Fiji [14]. For each case, the colocalization analyses were applied to each of the four to five images captured from each non-sequential sample.

### Movat pentachrome Staining

Tissue sections from the CM brain and skin specimens along with the non-CM brain and skin controls (**Table 1**) were sent for Movat Pentachrome staining to (iHisto, Salem, MA). MoticEasyScan Infinity 60 microscope was used for bright field scanning of each tissue section slide.

### Statistical analysis

Data are provided as mean ± standard error mean. Statistical packages in GraphPad Prism 9 were utilized for data normality and variance between the groups. Statistical differences were determined either by a two tailed t-test or One-way ANOVA (Brown-Forsythe and Welch ANOVA test) followed by Dunnett T3 post hoc multiple comparison test with significance considered as p<0.05.

## Results

### Sprouting endothelial cells in CM

It has been reported that CM blood vessels are predominantly enlarged and irregular [3, 4] and express angiopoietin-2, a potent angiogenic factor that can promote vascular remodeling[15]. Using fluorescently tagged Ulex europeus agglutinin-I (UEAI) to visualize human ECs in CM histological sections, we observed what appear to be actively sprouting ECs in numerous blood vessels (**Figure 1A-D**). We quantified the number of UEAI^pos^ blood vessels with at least one sprouting EC in brain and skin CM specimens as well as control brain and skin specimens for comparison (**Figure 1E**). Interestingly, there was a significant (p<0.0001) increase in sprouting UEAI^pos^ blood vessels in CM skin (38.2 ± 3.4%) compared to non-CM skin (9.8 ± 3.6%). In contrast, there was a trend but no significant (p=0.6628) increase in sprouting ECs in CM brain (9.9 ± 2.5%) compared to non-CM brain (4.5 ± 2.9%). There was a significant (p<0.0001) increase in blood vessels with sprouting ECs in CM skin compared to CM brain. This suggests a stronger angiogenic milieu may exist in the CM skin versus CM brain. Alternatively, features of the blood brain barrier may restrain sprouting behavior or the increased sprouting in the skin CM could be due to the increased *GNAQ* p. R183Q allelic frequency in the skin CM specimens versus brain CM specimens (**Table 1**) available for our study (**Supplemental Figure 1)**.

**Figure 1.**
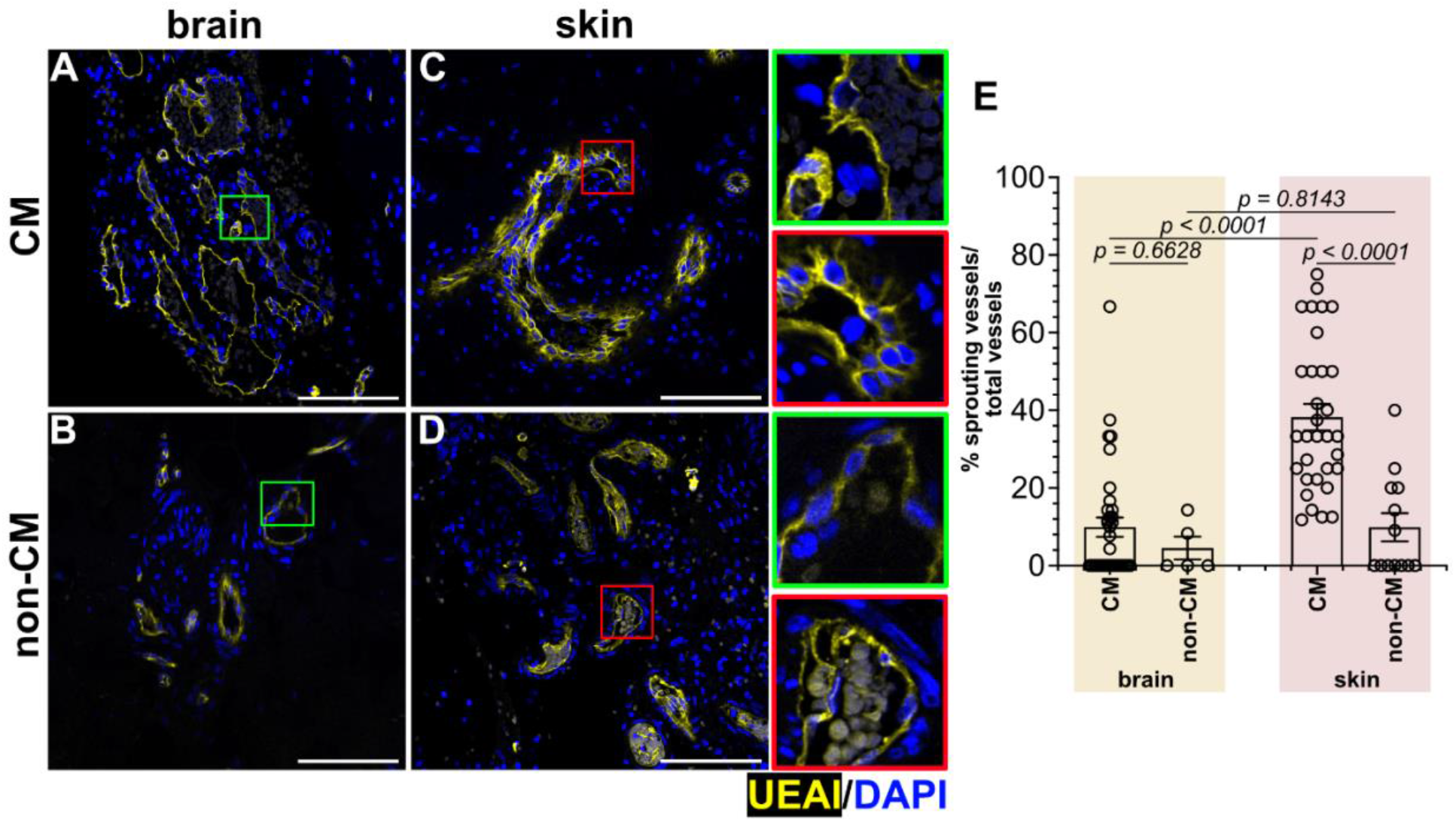
Ulex europaeus agglutinin I (UEAI) staining for endothelium in capillary malformation of brain and skin. UEAI (yellow), counter staining for nuclei by DAPI in (**A**) CM brain (n=4) (**B**) non-CM brain (n=2) (**C**) CM skin (n=4) (**D**) non-CM skin (n=2). High magnification images for CM brain (green box) and skin (red box) are presented as inserts. See Table 1 for information on tissue specimens. (**E**) Quantification of % sprouting vessels/total vessels. Scalebar = 100μm. The p-values were calculated by Brown-Forsythe and Welch ANOVA test followed by Dunnett T3 multi comparison test. 4-5 non-sequential sections were analyzed per tissue specimen.

### Extravascular fibrin in CM brain and skin specimens

We next looked at microenvironmental features. We performed Movat Pentachrome histological staining on the CM brain and skin specimens, and their respective controls to assess the extracellular matrix environment. Extracellular matrix is crucial for the maintenance of the vessel architecture as well as providing structure and function to the blood vessel. Previous Martius Scarlet Blue histological staining revealed extravascular fibrin in CM brain[4], which was confirmed here by Movat Pentachrome staining of CM brain specimens (n=4) (**Figure 2A**) and extended to reveal extravascular fibrin in CM skin specimens (n=4) (**Figure 2C**). In both brain and skin, extravascular fibrin was not detected in the controls (**Figure 2B, D**). These results uncover an apparent fibrin leakage in both brain and skin CM, suggesting comprised vascular integrity, in both brain and skin CM from patients.

**Figure 2.**
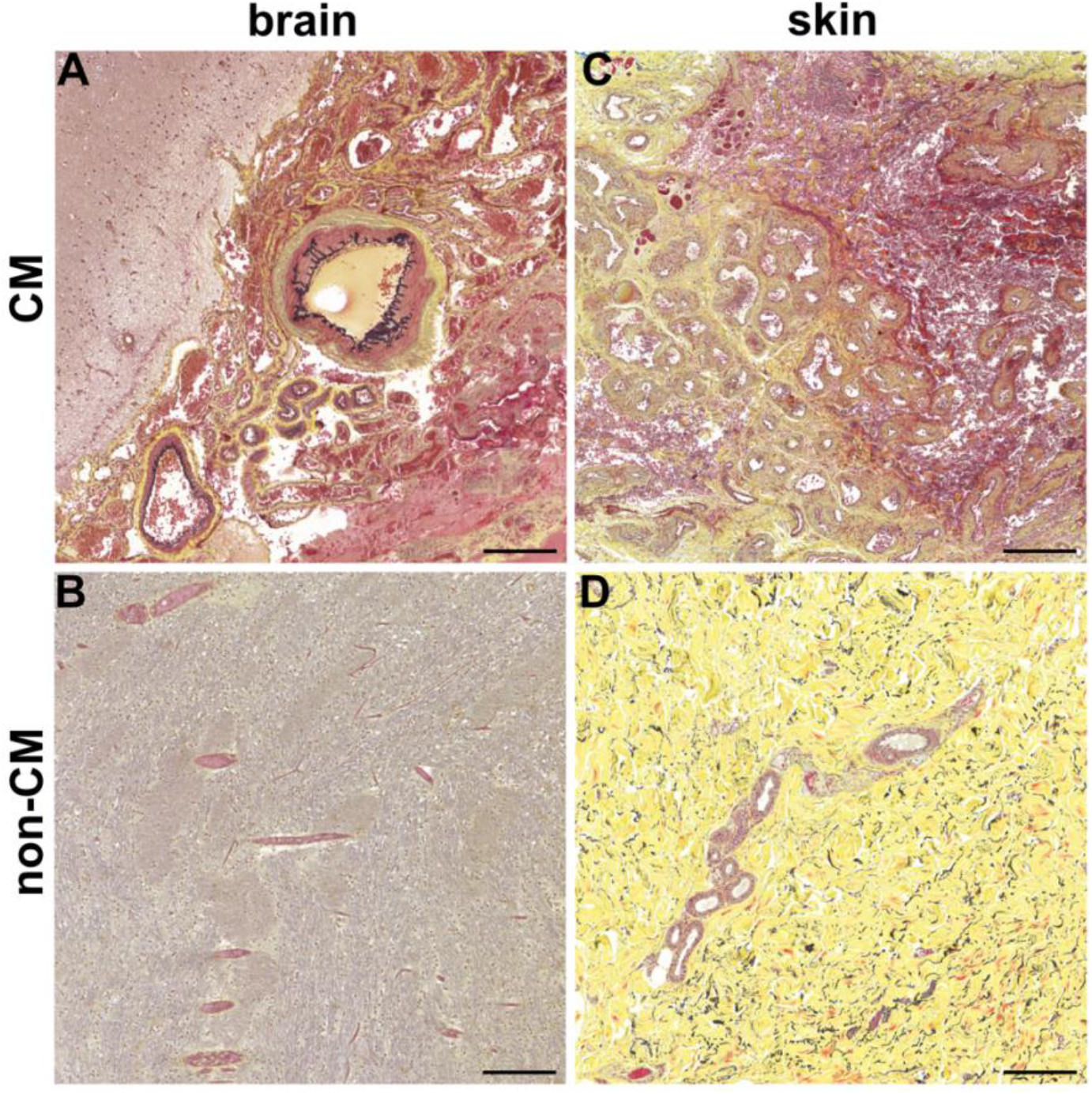
Movat pentachrome staining for elastin fibers and fibrin leakage in capillary malformation of brain and skin. (**A**) CM brain (n=4) (**B**) non-CM brain (n=2) (**C**) CM skin (n=4) (**D**) non-CM skin (n=2). Elastic Fibers (black); Collagen (yellow); Fibrin (bright red); Nuclei (blue/black). Scalebar = 200μm

### No detectable zona occludin1 in CM brain and skin specimens

Endothelial tight junctions are needed to create the critically important EC-EC barrier between the circulation and the abluminal environment. The fibrin leakage detected (**Figure 2**) and described previously[4] suggests a potential loss of EC-EC junctions. To assess this, we immunostained brain and skin CM sections for the tight junction protein zona occulin-1 (ZO1) and fluorescently tagged UEAI at the same time. Strikingly, we found majority of the blood vessels (UEAI^pos^) in the brain CM (**Figure 3A**) and skin CM (**Figure 3C**) were negative for anti-human ZO1 immunostaining. ZO1^pos^/UEA1^pos^ blood vessels were detected in control brain (**Figure 3B**) and skin (**Figure 3D**). The lack of ZO1 in CM blood vessels could be responsible for a diminished endothelial barrier and fibrin leakage.

**Figure 3.**
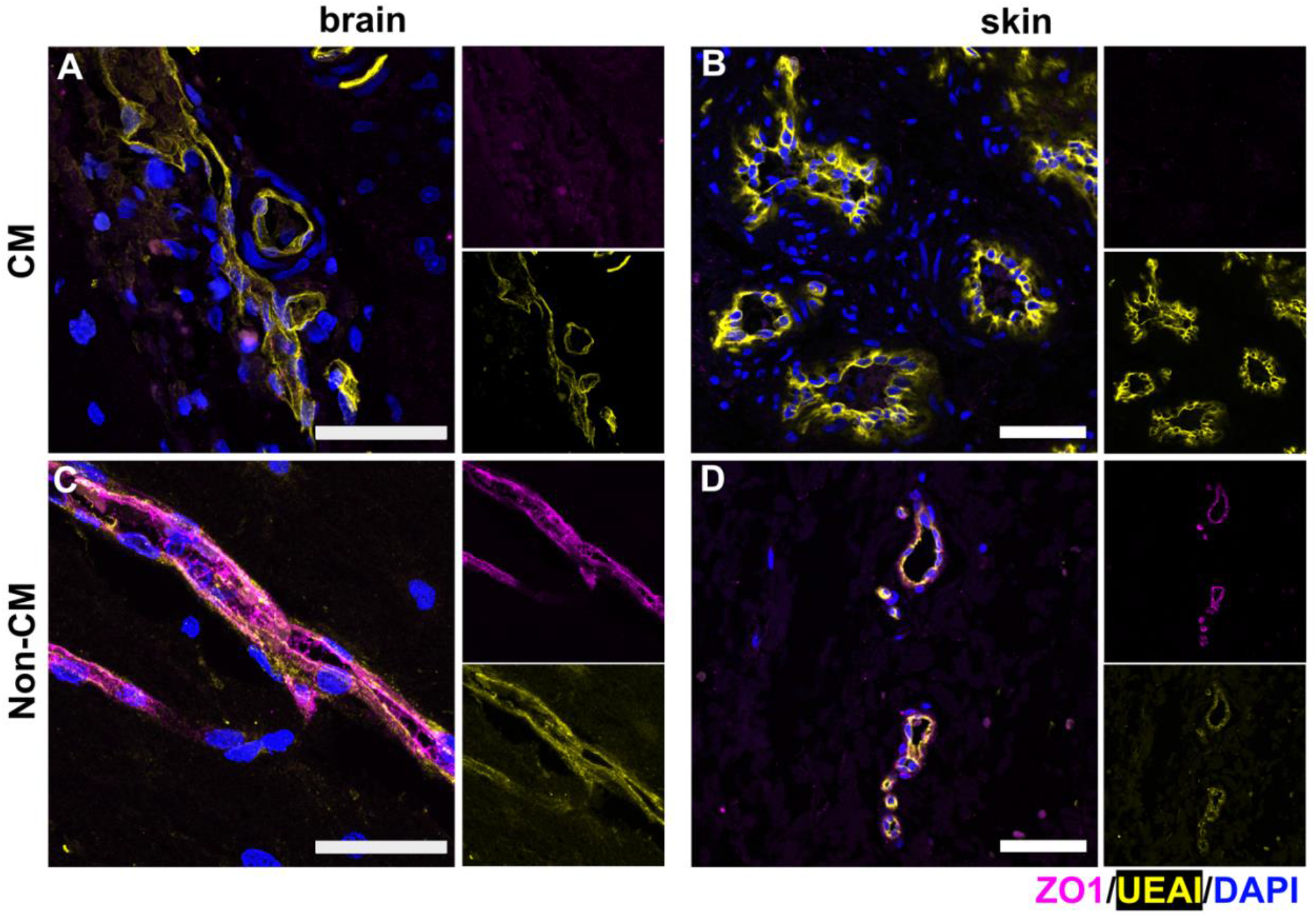
Loss of tight junction protein, zona occludin1 (ZO1), in capillary malformation of brain and skin. ZO1 (magenta), UEAI (yellow) and counter staining for nuclei by DAPI in (**A**) CM brain (n=4) (**B**) non-CM brain (n=2) (**C**) CM skin (n=4) (**D**) non-CM skin (n=2). Single channel images for UEAI (magenta) and ZO1 (yellow) are on the right of each merged image. Scalebar = 50 μm

### Disorganized mural cell coverage in CM skin compared to brain specimens

Mural cells are important in vascular development, vascular patterning, and vascular contractility and tone. Furthermore, mural cells induce endothelial quiescence and thereby dampen endothelial sprouting. We have recently shown that some of the enlarged vessels in the CM brain lack mural cells expressing the markers calponin, desmin, and alpha smooth muscle actin (αSMA)[4]. Herein, we compared these findings to CMs in the skin (**Figure 4 A, B**). As previously shown, enlarged vessels tend to lack mural cells, identified by reduced immunostaining for calponin, desmin and αSMA. A notable difference in mural cell coverage of CM vessels in skin specimens was that the mural cells were separated from the UEAI^pos^ endothelium. This was particularly striking for the calponin^pos^ and desmin^pos^ cells in CM skin. In contrast, αSMA^pos^ cells were detected on the abluminal side of the UEAI^pos^ endothelium suggesting heterogenous mural cell phenotypes in skin CM. By quantification, we found that 33% of the CM blood vessels in skin have mural cell coverage (magenta bar), ∼38% have no mural cell coverage (black bar) and ∼38% have a distant mural cell layer, labeled by anti-calponin or anti-desmin antibody (purple bar) (**Figure 4C**). The distance between the calponin^pos^ and desmin^pos^ cells and the UEAI^pos^ endothelium ranged from 5-15μm (**Figure 4D**) In contrast, control brain and skin specimens had a single layer of mural cell coverage surrounding the UEA1^pos^ endothelium (**Supplemental Figure 2**). Overall, CMs in the brain have fewer calponin^pos^ vessels compared to desmin and αSMA. Strikingly, CMs in the skin have distant mural cells labeled by anti-calponin and anti-desmin, which might be positive for αSMA.

**Figure 4.**
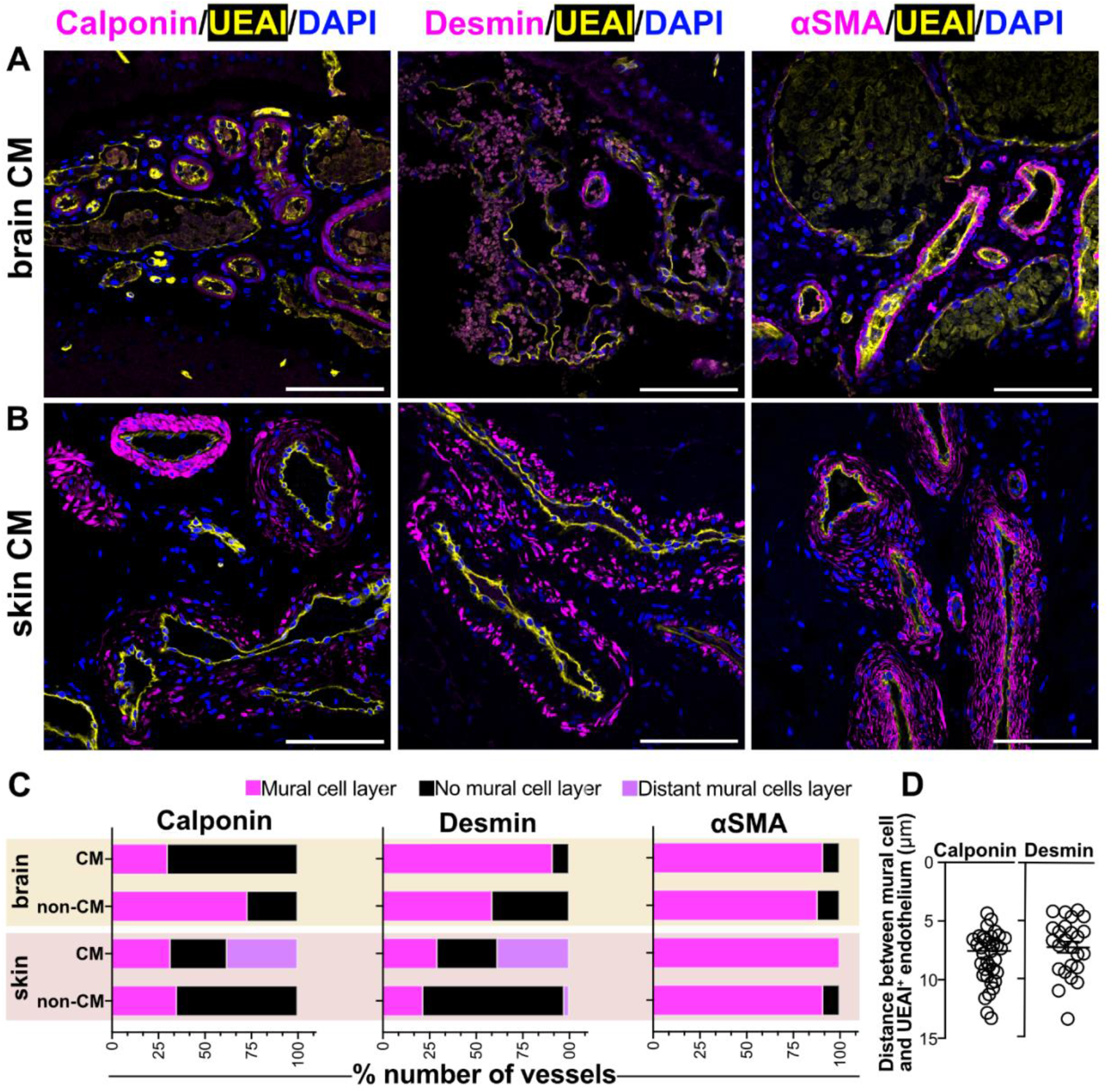
Mural cell environment in capillary malformation of brain and skin. Antibody staining for calponin (left), desmin (middle) and alpha-smooth muscle actin (αSMA) (magenta) and UEAI (yellow) and nuclei staining with DAPI (blue) in (**A**) CM brain (n=4) and (**B**) CM skin (n=4). Scalebar=100μm. See Table 1 for information on tissue specimens. (**C**) Quantification of % number of vessels with three distinct features: mural cell layer (magenta), no mural cell coverage (black) and distant mural cell layer (purple) in CM brain, non-CM brain, CM skin, non-CM skin. (**D**) Quantification of vessels with distance between the mural cell layer (Calponin (left) and Desmin(right)) and UEAI^pos^ endothelium (μm).

### Increased MRC1^pos^ and LYVE1^pos^ macrophages in brain and skin CM specimens

We showed previously there are significantly increased numbers MRC1^pos^/LYVE1^pos^/CD68^pos^ macrophages in the CM brain compared to non-CM brain controls[4]. We extended the analyses to CM in skin, analyzing CM brain in parallel (**Figure 5A, B, C, D**). We first looked if there are increased MRC1^pos^/LYVE1^pos^ macrophages in CM skin These cells are similarly elevated in skin CMs (p=0.0024) as in brain CMs (p=0.0310) compared to respective controls. (**Figure 5E**). When MRC1^pos^ cells were quantified, we found a significant (p<0.0001) increase in the number of MRC1^pos^ cells in CM brain compared to non-CM brain (**Figure 5F)**. In contrast, there was no difference (p=0.2592) in MRC1^pos^ cells in skin CMs compared to control skin. Hence, MRC1^pos^/LYVE1^pos^ macrophages are increased in both brain and skin CMs, but when a broader macrophage population was assessed, MRC1^pos^ cells, there was no significant increase in skin CMs compared to skin controls. Thus, while the perivascular macrophage component of CMs appears to be similar between brain and skin CMs, there are nuanced differences depending on the macrophage subset examined.

**Figure 5.**
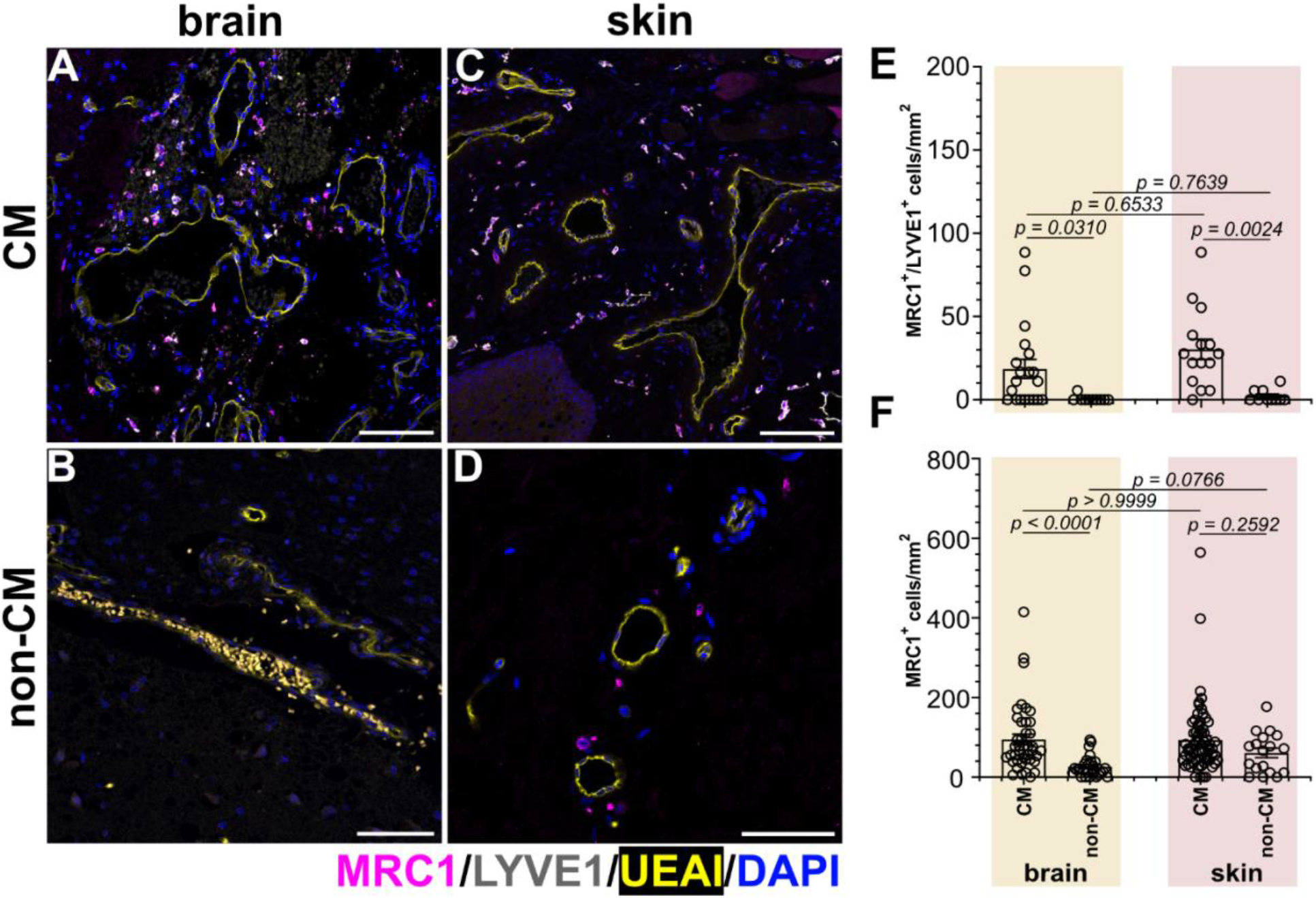
Mannose receptor C type 1 (MRC1^pos^) and lymphatic vessel endothelial hyaluronan receptor 1 (LYVE1^pos^) cells in capillary malformation of brain and skin. MRC1(magenta), LYVE1(grey), UEAI (yellow), and counterstaining for DAPI (blue) in (**A**) CM brain (n=4) (**B**) non-CM brain (n=2) (**C**) CM skin (n=4) (**D**) non-CM skin (n=2). See Table 1 for information on tissue specimens. (**E**) Quantification of colocalized MRC1^pos^ and LYVE1^pos^ cells in the CM brain and skin comparing to non-CM brain and skin. (**F**) Quantification of MRC1^pos^ cells in the CM brain and skin comparing to, non-CM brain and skin. The p-values were calculated by Brown-Forsythe and Welch ANOVA test followed by Dunnett T3 multi comparison test. 4-5 non-sequential sections were analyzed per tissue specimen. Scalebar = 200μm for panel A-C, 50μm for panel D.

## Discussion

In this study, we show CMs in brain and skin have similarities and differences in cellular and morphologic characteristics. Both consist of enlarged vessels with apparent fibrin leakage, perhaps due to lack of ZO1, and are surrounded by MRC1^pos^/LYVE1^pos^ macrophages. Notable differences were observed in skin CM. The endothelium of the skin CM vessels contained many ECs that appeared to be sprouting outward from the vessel lumen, which was much less apparent in brain CM. Furthermore, mural cells in skin CM appeared to be heterogenous and differentially localized compared to brain CM mural cells (**Figure 6**). Skin CM contained a ring of calponin^pos^/desmin^pos^ cells located 5-15μm away from the UEAI^pos^ endothelium. These histopathological characteristics – both the similarities and differences – are important to consider and may hold important clues for targeted molecular or tissue specific therapies for non-syndromic and syndromic CM.

**Figure 6.**
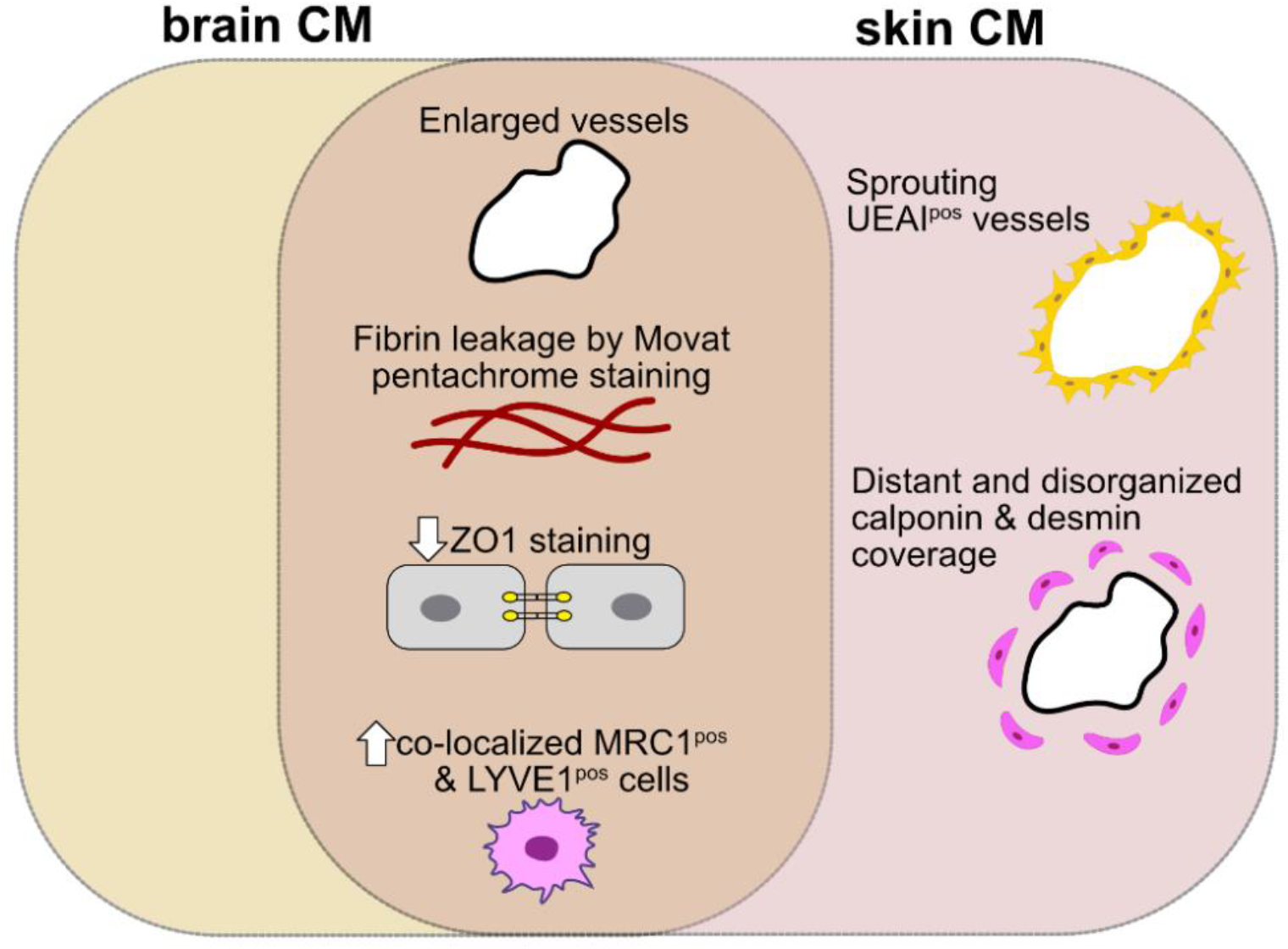
Summary comparing and contrasting brain and skin capillary malformation features by immunohistopathology.

The cellular composition of brain and skin is uniquely specialized based on the respective bio-mechano-physiological microenvironment. Skin has multiple layers, including epidermis (outer most layer serving as a protective barrier), dermis (middle layer that contains blood vessels, nerves, and hair follicles) and subcutaneous tissues (inner most layer). The skin contains various cell types such as keratinocytes, melanocytes, epithelial cells, fibroblasts, immune cells (such as macrophages and T cells) and nerve cells. In contrast, the brain has meninges covering and protecting the brain and spinal cord. The brain has a cerebrum, which is responsible for higher cognitive functions, cerebellum that regulates movement and balance, and brainstem controlling bodily functions such as heartbeat and breathing. The brain contains ECs, mural cells (pericytes and smooth muscle cells), neurons and glial cells (such as astrocytes, oligodendrocytes and microglia). The endothelium forms a blood brain barrier. All together the cells form a functional unit called “neurovascular unit”[16].

Brain and skin share a common embryological origin driving from the ectodermal layer during embryogenesis[17]. It remains unclear in the literature how the common ectodermal origin contributes to the specialized functions in the brain and skin in terms of bio-mechano-physiological characteristics. During murine embryonic vasculogenesis, mesodermal progenitor hemangioblasts migrate to the yolk sac giving rise to ECs and primitive blood cells. These same precursors within the embryo locally proliferate and undergo differentiation into a primary vascular plexus giving rise to the rest of the vasculature[18, 19]. The heterogeneity in ECs is linked to genetic factors, locality, soluble mediators, cell-to-cell contact, cell matrix interactions, pH, and mechanical forces (shear stress, strain and compression forces). In a healthy state, ECs are quiescent with turnover time over hundreds of days. However, during active angiogenesis, they can proliferate with a turnover time of less than 5 days[20]. In CM skin, we found UEAI^pos^ blood vessels to be sprouting in addition to undetectable ZO1 immunostaining, which we speculate are cells carrying the p.R183Q mutations[9, 10]. This remains an unanswered question due to lack of tools to distinguish mutant from non-mutant ECs in CM specimens. Future studies using CM specific animal models may address the role of Gα_q_ in ECs contribution to angiogenesis in a specific manner.

Interconnected endothelial junctions, namely, tight junctions, adherent junctions, and gap junctions are important for barrier function, tissue integrity, and cell-cell communication, respectively. Quiescent endothelium subjected to inflammation, immune cell extravasation, or angiogenic responses can dramatically impact vascular permeability and compromise the junctions. In the brain, the blood-brain barrier is maintained by a highly specialized organization of cellular environment. The physical barrier is maintained by tight junctions between ECs. It is well documented in the literature that the blood brain barrier is particularly enriched in tight junctions[21-23]. Emerging studies have also shown that there is significant crosstalk between tight junctions to regulate barrier function[24]. Moreover, during embryogenesis, ZO1 deficiency causes embryonic lethality by affecting yolk sac angiogenesis. This suggests that ZO1 may be functionally important for cellular remodeling and tissue organization in both embryonic and post-natal tissues[25]. As p.R183Q is a somatic activating mutation, it may cause an early deficiency and disruption of ZO1-containing tight junctions. This in turn could dysregulate early angiogenesis by failure to establish tight junctions and possibly other cell-cell connectors.

Mural cells (pericytes and smooth muscle cells) are found in most microvascular beds and their density and coverage depends on the tissue type[26]. Mural cells communicate with ECs through paracrine signaling by either diffusion or direct cell contact to allow for intercellular communication. Dysregulation in EC-mural cell communication can influence gene expression and thereby impact vascular permeability. Mural cells can arise from various progenitor cells depending on their origin, anatomical location and developmental time point. For example, in mice, neural crest derived mural cells give rise to choroid plexus[27] and thymus[28]. Recent studies have also shown that Col1a2 expressing-fibroblasts can give rise to pericytes[29, 30]. Mural cells are known to not only assist with the completion of vascular growth but to help establish correct blood flow patterns in the brain, also known as functional hyperemia[31, 32]. In the brain, mural cells also interact with astrocytes and neurons to establish and maintain the blood-brain barrier[33, 34]. In a pathological state, particularly in arteriovenous malformation, absence of pericytes surrounding the vessels has been reported[35-39]. In hereditary hemorrhagic telangiectasia type 2 patients with activin receptor-like kinase 1 loss of function, reduced mural cell coverage has been found with a potential mechanism linked to vascular endothelial growth factor[36]. In another study, Notch signaling has been linked to pericyte survival in arteriovenous malformations[40]. In CM brain and skin, we found many enlarged vessels devoid of mural cell coverage, Particularly in CM skin, we found mural cells separated from the endothelium by 5-15μM.

Macrophages are important for homeostasis and play diverse roles. They have the ability to sense pathogens and respond at the site of injury to begin repair processes. MRC1 – aka CD206 - expressing resident macrophages are known to facilitate phagocytosis. Moreover, it has been reported that there are subsets of macrophages that express LYVE1[41-43]. We previously showed that in the CM brain, there is an increased number of MRC1^pos^/LYVE1^pos^/ CD68^pos^ cells [4]. Herein, we show MRC1^pos^/LYVE1^pos^ cells are significantly higher in both CM brain and CM skin. Although it remains unknown where these perivascular macrophages originate from and the underlying recruitment mechanism in CMs, our previous study showed an increased adherence of macrophage cell lines (THP1 and U937) to *GNAQ* p.R183Q mutant ECs mediated by ICAM1[4]. As we found fibrin leakage in both brain and skin CMs, along with a lack of ZO1, one might speculate that the *GNAQ* p.R183Q endothelium preferentially allows macrophage adherence, and that this, coupled with the compromised tight junctions, may cause the vascular permeability.

There are limitations in our study. First, we relied on a small number of CM brain and skin samples as well as non-CM controls. The mutant/variant allelic frequency was higher in the skin CM specimens compared to brain CM specimens. Furthermore, the patients with skin CM were older than the patients with brain CM (**Table 1**). Despite the limited number of specimens, insights presented here comparing the tissue microenvironment are valuable and new. From the basic science perspective, it is important to take in account the tissue characteristics and the cellular environment to understand disease mechanisms as this might impact the potential drug target specific to the mutation driving the disease pathogenesis.

## Acknowledgements

The authors would like to thank the Vascular Biology Program Microscopy Imaging Core, and the Histology Lab at Boston Children’s Hospital. Research reported in this manuscript was supported by the National Heart, Lung, and Blood Institute, part of the National Institutes of Health, under Award Number 5R01HL127030. S.N. is supported by F32HL172637. C.B. was supported by Postdoc Mobility Fellowship from the Swiss National Foundation and by the Lisa’s Fellowship from the Sturge Weber Foundation. The content is solely the responsibility of the authors and does not necessarily represent the official views of the National Institutes of Health.

## Authors Contributions

SN conceived the project, collected data and material, designed, and executed experiments, and wrote the manuscript. JB supervised the project and wrote the manuscript. CB performed macrophage staining and quantification in neuropathology specimen. SA and AP provided the neuropathologic specimens. HK provided the cutaneous specimens.

---

**Supplementa1 Figure 1.**
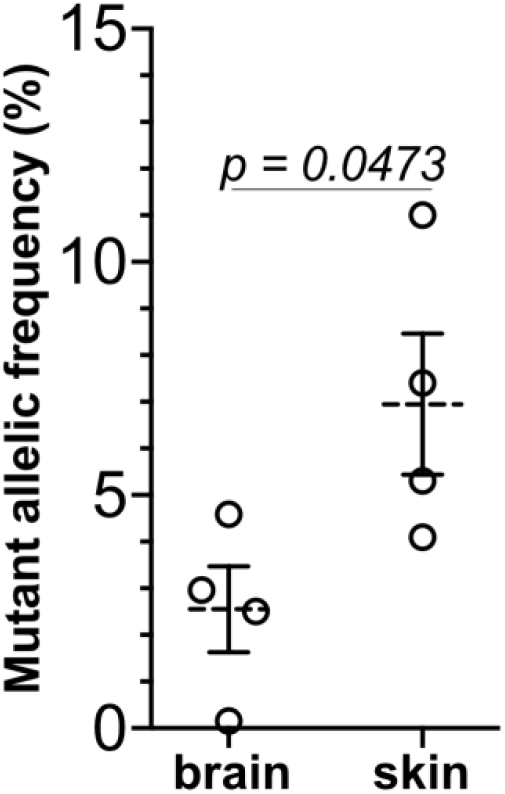
Mutant allelic frequency of brain vs. skin specimens (n=4). Two tailed t-test was performed. Normality of residuals was tested with Shapiro-Wilk (W) test. See Table 1 for information on tissue specimens.

**Supplementa1 Figure 2.**
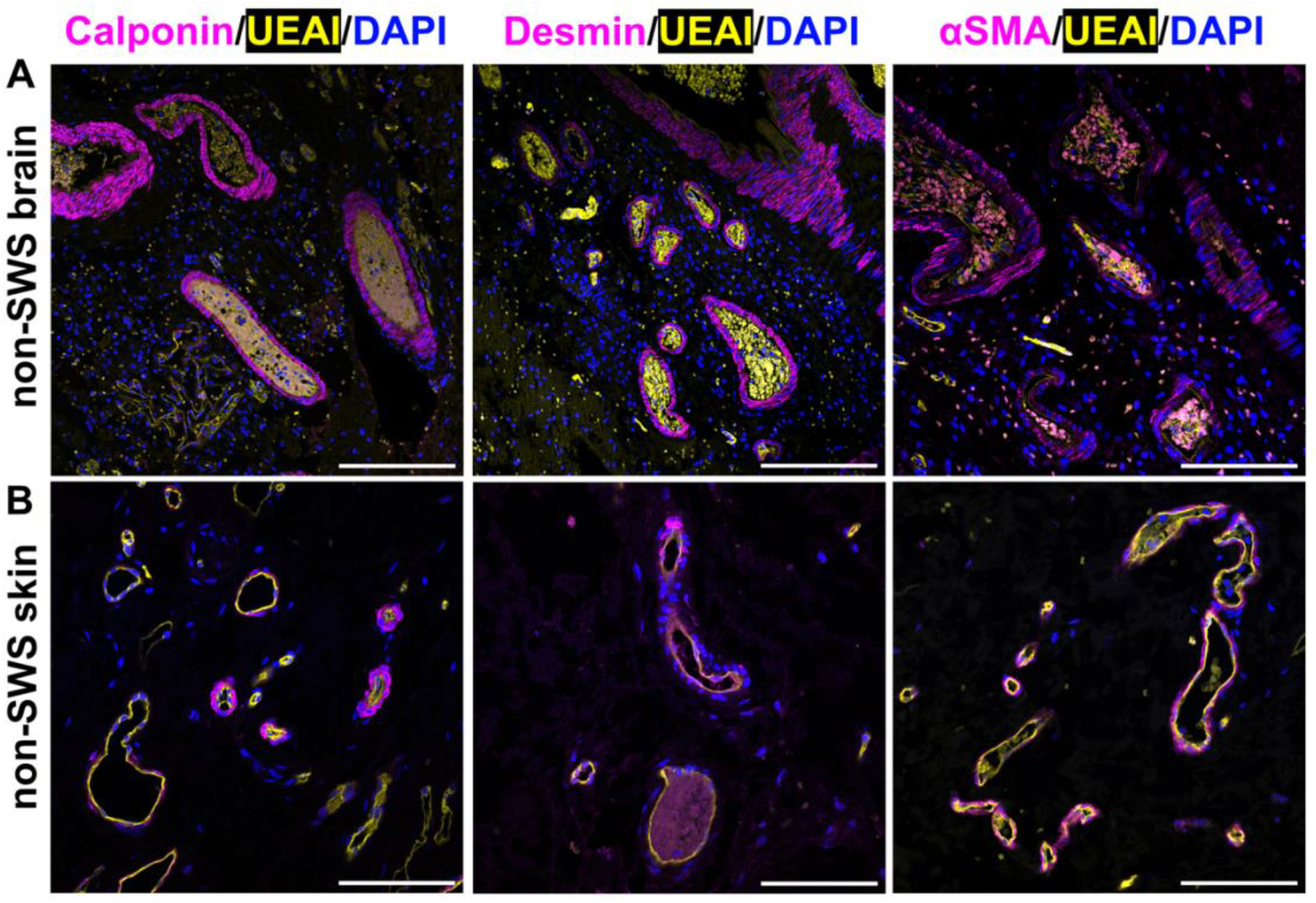
Mural cell environment in non-CM brain and skin. Antibody staining for calponin (left), desmin (middle) and alpha-smooth muscle actin (αSMA) (magenta) and UEAI (yellow) and nuclei staining with DAPI (blue) in (**A**) non-CM brain (n=2) and (**B**) non-CM skin (n=2). Scalebar=100μm. See Table 1 for information on tissue specimens.

**Table S1:**
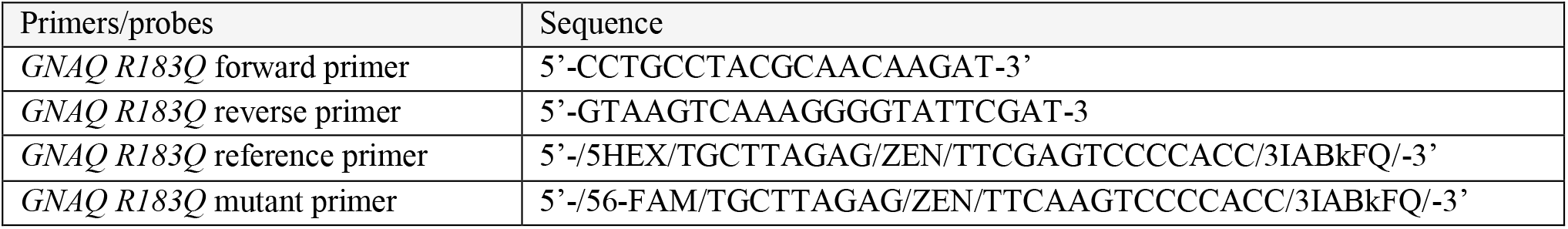
List of ddPCR primers and probes.

